# Von Willebrand Factor Deficiency Impairs Vascular Morphogenesis via Angiopoietin-2: Relevance for Gut Angiodysplasia

**DOI:** 10.1101/2025.09.09.675051

**Authors:** Adela Constantinescu-Bercu, Koval E. Smith, Shuo Yi Wong, Mattia Ballerini, Alessia Nastro, Benjamin G Wiggins, Daniela Pirri, Albert Li, Juun Evers, Olga Tsiamita, Matt Dibble, Charis Pericleous, Koralia Paschalaki, Graeme M. Birdsey, Michael A. Laffan, Suthesh Sivapalaratnam, Marco Rasponi, Anna M. Randi

## Abstract

Management of recurrent gastrointestinal (GI) bleeding is a clinical unmet need for patients with Von Willebrand disease (VWD) and is linked to the presence of gut vascular malformations (angiodysplasia). We previously demonstrated that von Willebrand factor (VWF) regulates angiogenesis and vascular integrity, the likely mechanism underlying angiodysplasia. VWF controls the storage of the angiogenesis regulator Angiopoietin-2 (Angpt-2) in endothelial cells (EC), suggesting a candidate for the genesis of angiodysplasia; however, no direct evidence of the role of Angpt-2 in VWF-dependent angiogenesis is available. Here we use VWF-deficient HUVEC, and endothelial colony forming cells (ECFCs) from severe VWD patients and find that loss of VWF in EC results in increased Angpt-2 expression through a positive feedback loop via the Angpt-2-TIE2-AKT-FOXO1 pathway. We also show an imbalance of the Angpt/Tie2 pathway *in vivo.* In the gut of VWF-deficient mice, Angpt-2 expression is increased whilst Angpt-1 expression is decreased; this correlates with reduced expression of the pericyte marker NG2. These data suggest that VWF regulates the Angpt/Tie2 balance in the gut. To investigate the functional defects caused by loss of VWF, we use a fibrin bead assay and show that VWF-deficient HUVEC present increased sprouting. We develop a microfluidic model of 3D vasculogenesis/angiogenesis and find that ECFCs from VWD patients exhibit defective remodeling and abnormal lumen formation compared to healthy controls. Importantly, inhibition of Angpt-2 reduces sprouting in VWF-deficient HUVEC and normalises vascular networks in ECFCs from severe VWD, suggesting Angpt-2 inhibitors may be effective in VWD patients with GI bleeding and angiodysplasia.

**Key points:**

1. Endothelial VWF regulates multiple steps of angiogenesis, including sprouting and lumen formation.
2. VWF regulates Angpt-2 storage and expression, and Angpt-2 blockade normalises defective angiogenesis in VWD ECFCs.

**Visual Abstract:** VWF deficiency in HUVECs results in increased Angpt-2 release and expression via the Tie2-Akt-FOXO1 pathway.
**A.** Model pathway for the regulation of Angpt-2 levels by VWF. Loss of VWF results in increased Angpt-2 release from endothelial cells; Angpt-2 binds to and inhibits the Tie2 receptor, decreasing its phosphorylation as well as the downstream phosphorylation of Akt and FOXO1; this leads to FOXO1 activation and an increase expression of Angpt-2, generating a feedback loop. **B.** Model for the pathogenesis of angiodysplasia in VWD. In the absence of VWF, the increase in Angpt-2 disrupts vascular morphogenesis through multiple mechanisms: increased sprouting, impaired vascular remodelling and an imbalance between Angpt-1 and Angpt-2 in the gut. Figure generated with Biorender.

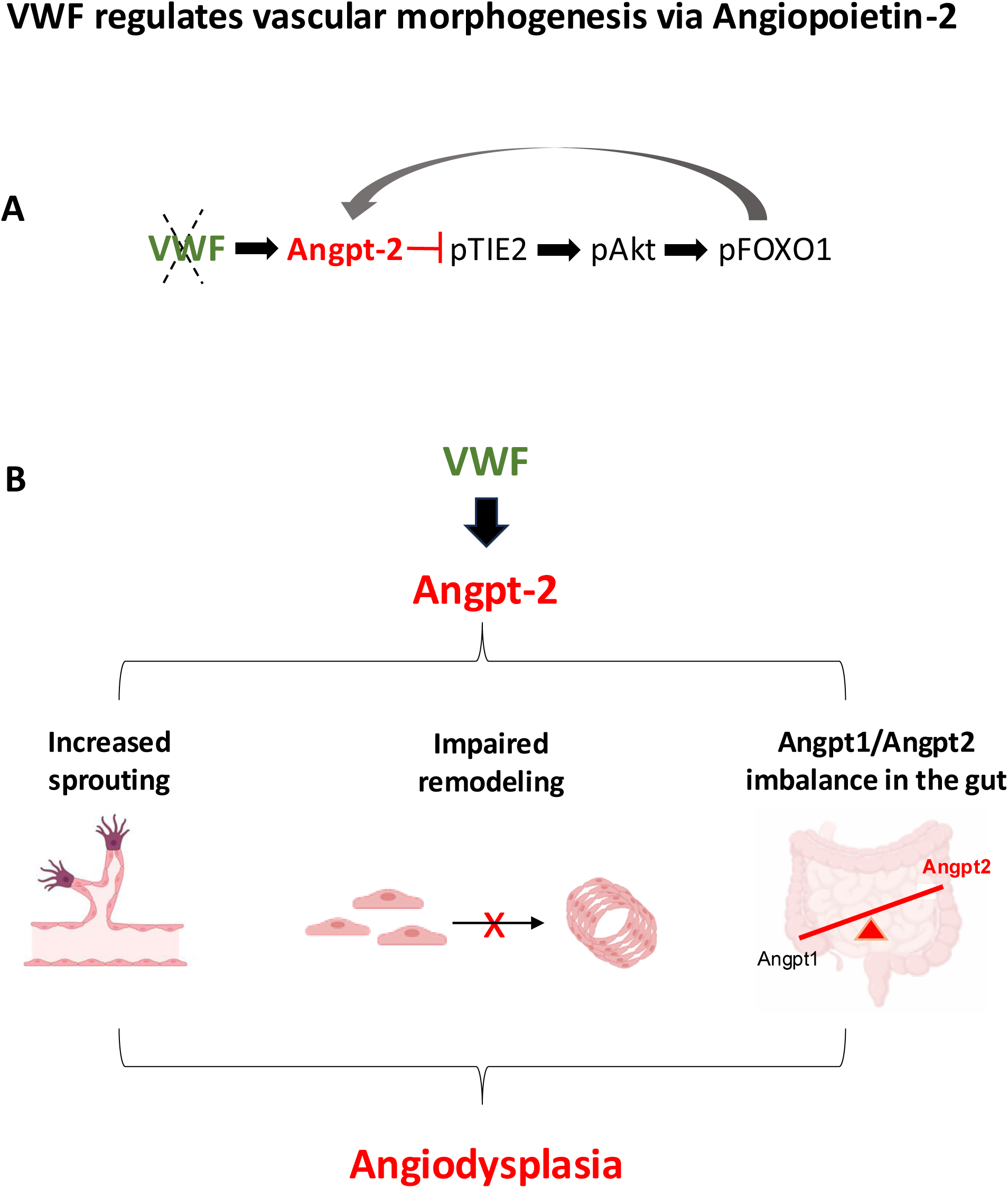

## Introduction

Von Willebrand factor (VWF) is a large glycoprotein best known for its roles in haemostasis^1, 2^. Congenital decrease or dysfunction of VWF lead to the most common human inherited bleeding disorder, Von Willebrand disease (VWD)^3^. In a subset of patients with congenital VWD, vascular malformations in the gastrointestinal (GI) tract (angiodysplasia) are present; these are characterized by fragile, leaky mucosal vessels, and can lead to severe, intractable bleeding^4–6^. The prevalence of angiodysplasia in VWD varies according to reports, between 5 and 20%^5, 7, 8^, and is more prevalent in patients with very low VWF levels or lacking VWF high molecular weight (MW) multimers^8–10^. Angiodysplasia is also common in acquired forms of VWD (also called acquired von Willebrand syndrome), associated with conditions such as aortic stenosis, paraproteinemia and left assisted ventricular device (LVAD)^11–14^. The condition can be clinically challenging, with severe, recurrent GI bleeding often refractory to VWF replacement therapy^15, 16^.

Angiodysplasia is thought to be associated with disrupted angiogenesis, but the mechanisms are unknown. Our previous work identified a role for VWF in angiogenesis and vascular development, suggesting a possible pathogenetic mechanism for angiodysplasia in VWD. *In vitro* and *in vivo* studies^17, 18^. suggested that VWF may deliver inhibitory signals during angiogenesis, possibly promoting maturation. However, in a model of wound healing, VWF-deficient mice showed decreased angiogenesis^19^. This complex picture indicates that VWF may exert different effects on angiogenesis depending on tissue and microenvironment.

The mechanisms through which VWF controls blood vessel formation are still unclear. VWF is present in 3 pools: cells (EC and platelets), plasma and the sub-endothelium. VWF monomers synthesized by EC are processed to assemble into multimers which reach a MW up to 20,000kDa. These are packaged into Weibel-Palade bodies (WPB) for basal and regulated secretion^20, 21^. VWF is essential for the formation of WPB, which contain proteins with diverse vascular roles^22^. One of these is Angiopoietin (Angpt)-2^23^, a growth factor part of the Angpt/Tie-2 signaling pathway, which regulates vascular homeostasis, angiogenesis and vascular integrity^24^. In most circumstances, Angpt-2 competes with its homologue Angpt-1 for binding to the endothelial receptor Tie2, synergizing with VEGF to promote angiogenesis^24^. Angpt-2 is synthesised by EC and stored in WPB, whilst Angpt-1 is mainly produced by mural cells. In healthy endothelium, levels of Angpt-2 are low; its expression is upregulated during inflammation and angiogenesis^25^.

We have previously shown that inhibition of VWF expression in EC *in vitro* causes increased release of Angpt-2, due to loss of WPB^17^. Here we show that VWF also controls Angpt-2 synthesis in EC via a positive feedback loop through the Angpt-2-TIE2-AKT-FOXO1 pathway; we confirm disruption of this pathway in the gut of VWF-deficient mice. Finally, we explore the cellular consequences of VWF loss in a new microfluidic vasculogenesis/angiogenesis model. This reveals abnormal vascular remodelling and defective lumen formation in ECFCs from patients with severe VWD and angiodysplasia. Crucially, both can be normalized by the anti-Angpt-2 blocking antibody MEDI3617. This study validates the Angpt-2 pathway as a promising candidate for therapeutic approaches in patients with VWD and provides new insight into the cellular angiogenic defects in VWD ECFC, which may lead to novel therapeutic targets.

## METHODS

### Cells

Human umbilical vein endothelial cells (HUVEC, Lonza) were cultured in endothelial cell growth medium 2 (EGM-2, Lonza). Endothelial colony forming cells (ECFCs) were isolated from healthy controls (HCs) or VWD type 3 patients as described^17, 26^, following informed consent in accordance with the Declaration of Helsinki. EGM-2 was supplemented with 20% human serum (298189, Merck Life Sciences). More details provided in Supplemental Methods. Human dermal skin fibroblasts (Detroit 551, Public Health England) were purchased from the European Collection of Cell Cultures (ECACC), maintained in M199 supplemented with 10% FBS at 37°C and 5% CO2, and used between passage 2-4.

### siRNA treatment of endothelial cells

Two different siRNA sequences were used to target VWF as described^17^. Expression of Forkhead Box O1 (FOXO1) was inhibited using pre-validated siRNA (Qiagen)^27^ (details in Supplemental Methods).

### Mice breeding and tissue isolation

Experiments on global VWF-deficient mice^28^ (KO mice) (The Jackson Laboratory) or littermate controls (WT mice) were conducted according to Imperial College London-approved protocols, in compliance with the UK Animals (Scientific Procedures) Act of 1986. Both male and female mice (6–12 weeks old) were used. Small intestine sections (duodenum, jejunum and ileum) were isolated from WT and KO mice. 1cm of each segment was snap frozen for RNA extraction. Tissues were homogenised in Trizol using Precellys Lysis Kit (Thermo Fisher) according to manufacturer’s instructions. Homogenized tissues were mixed with chloroform and spun for 15min to separate the aqueous phase and processed for RNA extraction.

### Real-time polymerase chain reaction

Total RNA from HUVEC, ECFCs and mouse gut tissue was isolated using the RNeasy kit (Qiagen) and reverse transcribed into cDNA using Superscript III Reverse Transcriptase (Invitrogen). Quantitative real-time PCR was performed using PerfeCTa SYBR Green Fastmix (Quanta Biosciences) on a Bio-Rad CFX96 system. Gene expression values were normalized to *GAPDH* expression.

### ELISA

Supernatants were collected and processed as described^17^; VWF ELISA was performed as described^17^. Human Angpt-2 ELISA (DY623, R&D Systems) were performed according to manufacturers’ instructions. ELISAs were performed on supernatants and lysates; results were normalised to the total amount of protein in the lysate samples.

### Immunoblotting

Cell lysates were immunoblotted using antibodies to: VWF (Dako), phospho-Akt Ser-473, pan-Akt, phospho-FOXO1 Thr-24, FOXO1, Tie2 (Cell Signalling). Quantification was performed by densitometry and normalized against GAPDH (Millipore). Details are provided in Supplemental Methods.

### Fibrin Bead Assay

HUVEC were treated with siCTL or siVWF 24 hours prior to coating onto cytodex 3 beads (GE Healthcare) as described^29, 30^ (details in Supplemental Methods). 20,000 fibroblasts were seeded on top of the fibrin clot in EGM-2 supplemented with 2%FCS, IgG control or Angpt-2 IgG neutralising antibody (AZ 3.19.3)^31^ at 5nM. For experiments using Tie-2-Fc, 2.5ng/ml were added to complete EGM-2 or PBS. Images of beads were captured at day 5 on Olympus IX70 microscope using 10X objective and analysed using ImageJ. The number of sprouts/bead was determined; a minimum of 30 beads/condition was analysed.

### Microfluidic device fabrication

The microfluidic devices were produced through soft lithography using the design recently published in^32^, with modifications. The cell culture layer of the microfluidic device was fabricated using replica moulding with polydimethylsiloxane (PDMS, Sylgard 184, Dow Corning). Briefly, PDMS, mixed with the curing agent in a 10:1 ratio, was poured into the moulds and cured at 65°C for 2 hours to allow crosslinking. Layers were irreversibly bonded to glass coverslips through air plasma treatment (Harrick Plasma, 25W, 50s), or oxygen plasma (Diener Electronic, 45W, 35s, 0.3mBar O₂).

### 3D vasculogenesis/angiogenesis assay

HUVEC or ECFCs, resuspended at 10million cells/ml, were embedded in a fibrin gel, consisting of thrombin (Sigma, 1.5U/ml), fibrinogen (Sigma, 6mg/ml) and EC in a 2:2:1 ratio, and injected in the central channel of the microfluidic devices. After 5 minutes of incubation at 37°C for fibrin polymerization, the devices were hydrated with EGM-2 supplemented with 2%aminocaproic acid (ACA), in the presence or absence of the blocking anti-Angpt-2 IgG antibody (MEDI3617, Bio-Techne, 10nM) or a human IgG isotype control (Bio-Techne). The antibody-containing medium was replenished every 24 hours. Angiogenic networks were monitored using brightfield microscopy at 2, 4, 24, 48 hours. Image analysis was performed using ImageJ Angiogenesis analyser. Total network length was quantified.

### Immunofluorescence: Endothelial cells in 2D

HUVEC or ECFC were immunostained as described^33^ using antibodies to VWF (Dako), VE-cadherin & Angpt-2 (Santa Cruz Biotechnology), VE-cadherin (BD-Biosciences), FOXO1 (Cell Signalling), p-Tie2 (R&D Systems) and DAPI (Invitrogen). Further details in Supplemental Methods.

### Immunofluorescence: Angiogenic networks in 3D

Angiogenic networks in the microfluidic devices were fixed after 48 hours with 4%PFA, then blocked and permeabilized with 3%BSA, 0.25%Triton-X100, 5%FBS in PBS. Networks were stained using antibodies to VWF and VE-cadherin (as above). At least 3 images/microfluidic device were acquired using a Leica Stellaris 8 confocal microscope and analysed using ImageJ (%area covered by the networks) or Imaris Analysis Software (Oxford Instruments) (3D-rendering to quantify angiogenic network volume).

### Data Analysis

Data is presented as mean ± standard deviation (SD) of at least three independent experiments. Statistical analysis was performed using GraphPad Prism software 10.0.

## RESULTS

### VWF controls Angpt-2 expression through a Tie2-Akt-FOXO1 dependent pathway

We have previously shown that VWF-deficiency causes increased release of Angpt-2 from EC^17^ (Figure 1A). Accordingly, intracellular Angpt-2 levels were decreased in VWF-deficient HUVEC (Supplemental Figure 1A). Angpt-2 acts as a context dependent agonist/antagonist of Tie2^34^; we speculated that in VWF-deficient cells, increased Angpt-2 release could cause a decrease in Tie2 phosphorylation, as reported^35^. Indeed, Tie2 phosphorylation was significantly decreased in VWF-deficient cells (Figure 1B and Supplemental Figure 1B).

**Figure 1.**
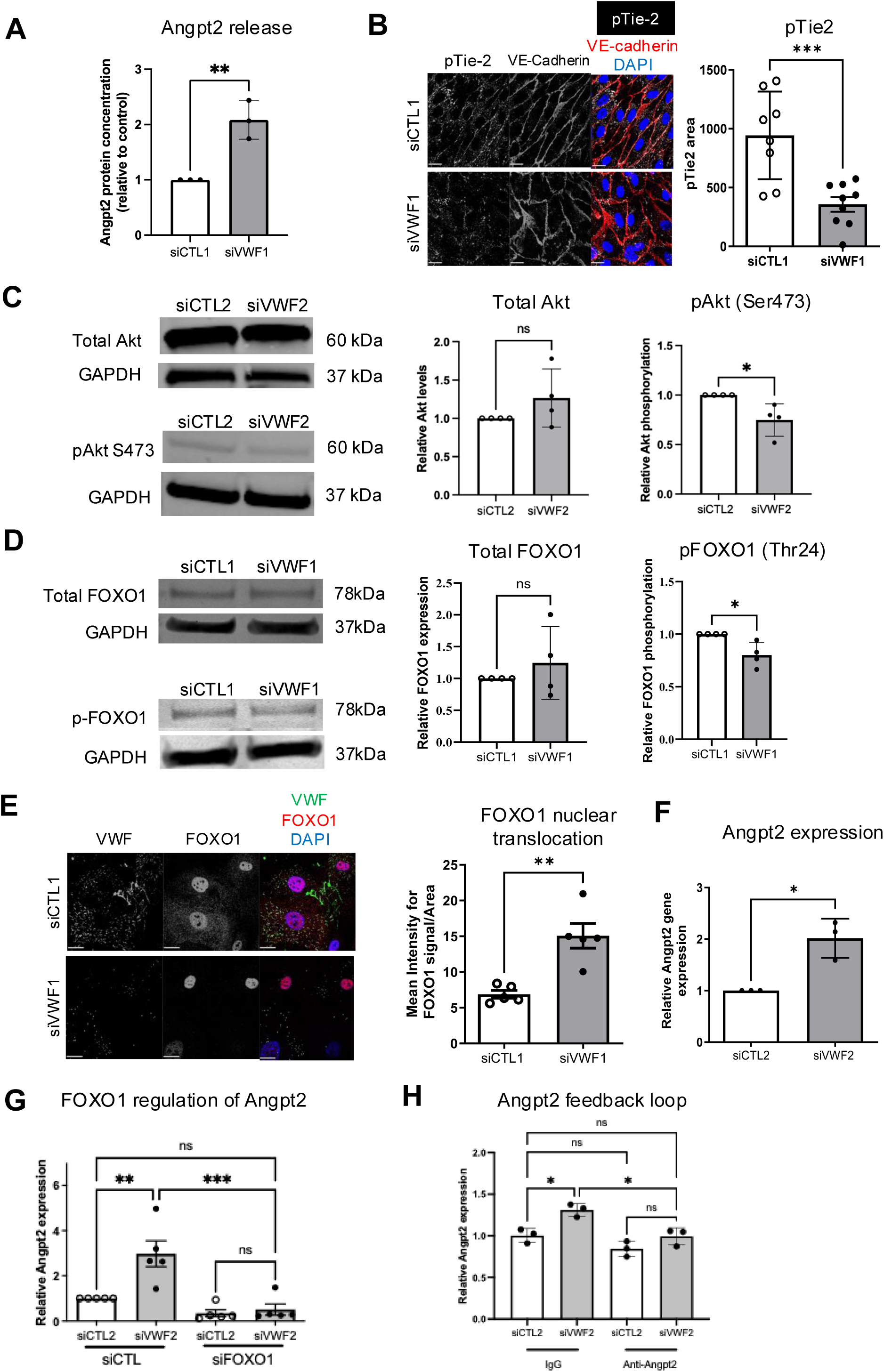
VWF deficiency in HUVECs results in increased Angpt-2 release and expression via the Tie2-Akt-FOXO1 pathway. **A.** Increased Angpt-2 release following inhibition of VWF expression using siVWF1. n=3. ****p<0.0001, Student’s t-test. **B.** Representative images of HUVECs transfected with siCTL1 or siVWF1, fixed after 48 hours and stained for VE-cadherin (red), pTie2 (white) and DAPI (blue). pTie2 stained area was quantified. n=8 images/condition from 3 different transfections. ***p<0.001, Student’s t-test. Scale bar 50μm. **C.** Western blot of pAkt (S473) and total Akt in siCTL2- and siVWF2-transfected HUVECs. n=4 lysates/condition, from 2 separate transfections. **p<0.01, unpaired Student’s t-test. **D.** Western blot and quantification of total FOXO1 and pFOXO1 in siCTL1 and siVWF1-treated HUVECs. n=4 different transfections. *p<0.05, unpaired Student’s t-test. **E.** Representative images of siCTL1 and siVWF1-treated HUVECs confirming decreased VWF levels (green) following siVWF1 treatment and increased translocation of FOXO1 (red) to the nucleus (blue). Scale bar 50μm. Data was quantified by analysing the ratio between mean intensity of signal and area of FOXO1 staining. **p<0.001, unpaired Student’s t-test. **F.** Angpt-2 expression in the presence of siCTL2/siVWF2. n=3 different transfections. *p<0.05, unpaired Student’s t-test. **G.** Angpt-2 expression in HUVECs transfected with siCTL2 or siVWF2, as well as siCTL or siFOXO1. n=3-5 lysates/condition. *p<0.05, **p<0.01, ***p<0.001, One-way ANOVA with multiple comparisons. **H.** Angpt-2 expression in siCTL2 and siVWF2-transfected HUVECs, in the presence of control IgG or the blocking antibody against Angpt-2 (MEDI3617). *p<0.05, One-way ANOVA with multiple comparisons. All data are plotted as mean +/- S.D.

Tie2 controls phosphorylation of the serine/threonine kinase AKT^36^ and the downstream transcription factor FOXO1. Phosphorylation of FOXO1 leads to its cytoplasmic retention, thereby inhibiting its transcriptional activity. Conversely, inhibition of FOXO1 phosphorylation leads to its nuclear translocation and increased transcriptional activity^37^. We hypothesized that decreased Tie2 phosphorylation in VWF-deficient cells could suppress the AKT-FOXO1 axis. Indeed, we observed a significant reduction in Akt phosphorylation (Ser473) (Figure 1C), FOXO1 phosphorylation (Thr24) (Figure 1D) and a significant increase in FOXO1 nuclear translocation in VWF-deficient HUVEC compared to controls (Figure 1E), suggesting increased FOXO1 activity.

FOXO1 drives Angpt-2 expression in EC^27, 38, 39^. In line with increased FOXO1 nuclear translocation, Angpt-2 mRNA levels were significantly increased in VWF-deficient cells compared to controls (Figure 1F and Supplemental Figure 1C), as demonstrated with two siVWF sequences (Supplemental Figure 1D-E). Angpt-2 mRNA stability was similar in VWF-deficient cells, in line with transcriptional regulation (Supplemental Figure 1F-G). Crucially, inhibition of FOXO1 expression decreased Angpt-2 mRNA expression in VWF-deficient cells (Figure 1G and Supplemental Figure 1H). These results demonstrate that loss of endothelial VWF causes an increase in Angpt-2 expression via the transcription factor FOXO1.

These data suggest a feedback loop regulating Angpt-2 expression in VWF-deficient cells, dependent on loss of WPB which causes Angpt-2 release and subsequent activation of the Tie2-AKT-FOXO1 pathway, which drives further Angpt-2 expression (Visual Abstract A). To test this model, we treated VWF-deficient HUVEC with the anti-Angpt-2 blocking antibody AZ3.19.3 or control IgG. Blockade of Angpt-2 significantly reduced Angpt-2 mRNA expression in VWF-deficient HUVEC (Figure 1H and Supplemental Figure 1I), confirming the presence of a positive feedback loop regulating Angpt-2 expression in VWF-deficient HUVEC.

### VWF^-/-^ mice present an Angpt-1/Angpt-2 imbalance in the gut

To investigate whether VWF regulates Angpt-2 expression in the gut, Angpt-2 mRNA levels were measured in whole tissue samples from small intestine regions (duodenum, ileum and jejunum) of VWF-deficient mice and wild-type littermates (Figure 2A). Analysis showed a significant increase in Angpt-2 expression levels in the jejunum of VWF-KO mice (Figure 2B and Supplemental Figure 2A). Importantly, levels of endothelial and epithelial markers (CD31, eNOS and CDH5) were not significantly different between WT and KO jejunum (Supplemental Figure 2D), indicating no major difference in cellular composition. Given that Angpt-2 influences pericyte coverage^40–42^, we investigated expression of the pericyte marker NG2 and of Angpt-1, which is produced by pericytes. Our results revealed a significant decrease in NG2 and Angpt-1 expression in the small intestine overall, mainly driven by the jejunum area (Figure 2C-F and Supplemental Figure 2B-C). A positive correlation between NG2 and Angpt-1 expression overall and in the jejunum alone (Supplemental Figure 2E-F) suggests that the decrease in Angpt-1 is driven by decreased pericyte coverage in the gut of VWF-KO mice. These data indicate that lack of VWF causes an imbalance in the Angpt/Tie2 pathway in the gut (Visual Abstract B).

**Figure 2.**
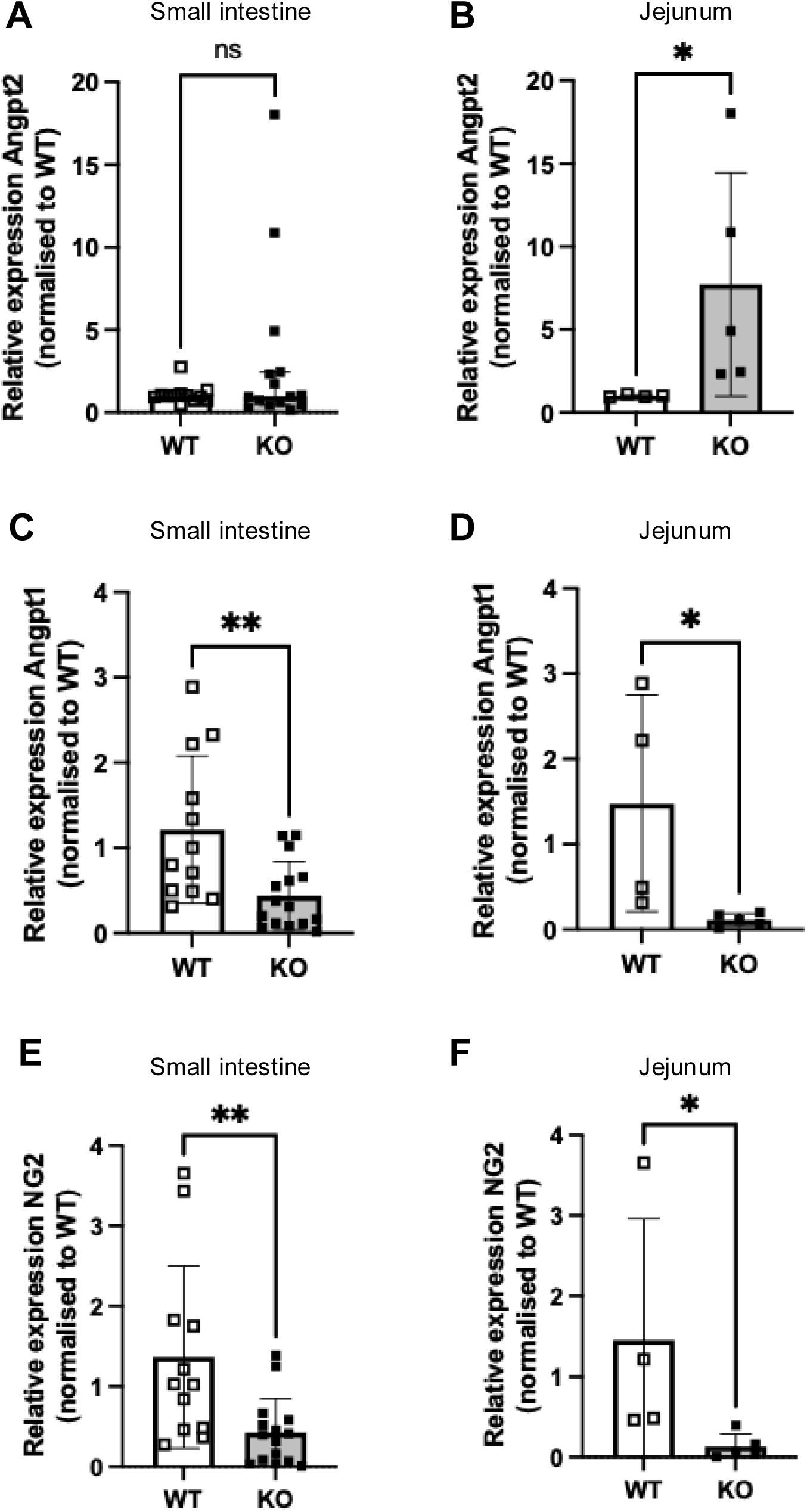
VWF^-/-^ mice show an imbalance in Angpt-2 / Angpt-1 expression in the gut (jejunum). **A.** Angpt-2 mRNA expression in the small intestine overall. **B.** Angpt-2 mRNA expression in the jejunum. **C.** Angpt-1 mRNA expression in the small intestine overall. **D.** Angpt-1 mRNA expression in the jejunum. **E.** NG2 mRNA expression in the small intestine overall **F.** NG2 mRNA expression in the jejunum. n=4 WT, n=5 KO. All data normalized to WT, *p<0.05, **p<0.01, Mann-Whitney test. All data are plotted as mean +/- S.D.

### Blocking Angpt2 normalises VWF-dependent increase in sprouting angiogenesis

Since our observation that VWF controls Angpt-2 storage in EC^17^, the hypothesis that Angpt-2 mediates VWF-dependent control of angiogenesis has been widely discussed. However, experimental evidence in support of this hypothesis is missing. To determine whether VWF regulates sprouting angiogenesis through the Angpt-2/Tie-2 pathway, we used the *in vitro* fibrin bead assay, a co-culture model where fibroblasts provide factors that promote endothelial sprouting^30, 43^. HUVEC were treated with control or VWF siRNA for 24 hours (Supplementary Figure 1B). Inhibition of VWF expression induced a significant increase in the number of sprouts compared to siCTL-treated cells (Figure 3A-B), indicating that VWF affects sprouting angiogenesis. Tie2-Fc significantly reduced enhanced EC sprouting in VWF-deficient cells (Figure 3A), to levels comparable to siCTL cells. These data indicate that the Angpt/Tie-2 pathway mediates the enhanced endothelial sprouting in siVWF-treated HUVEC.

**Figure 3.**
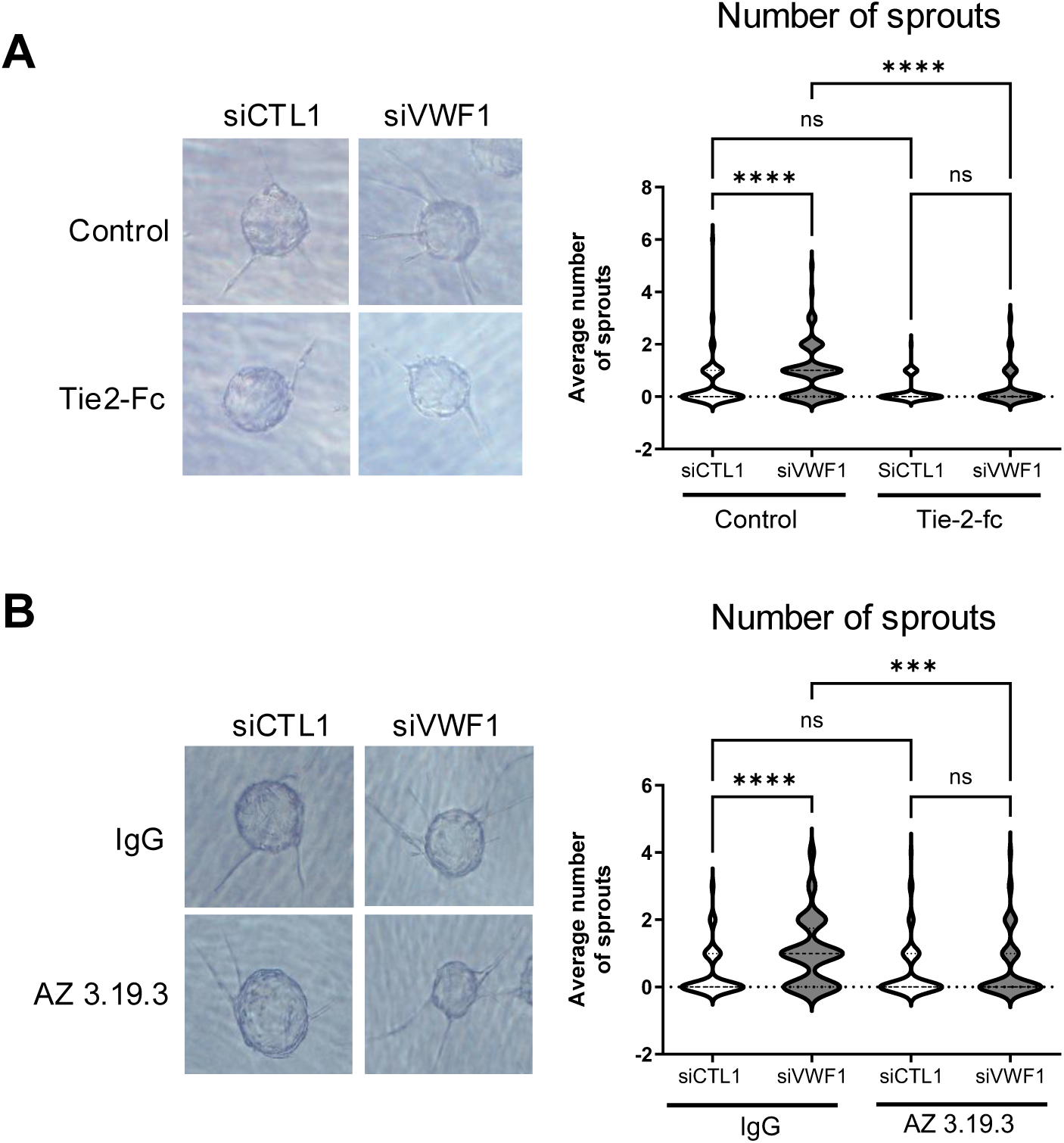
VWF regulates sprouting angiogenesis via Angpt-2. **A.** Fibrin bead sprouting angiogenesis assay in the presence or absence of Tie2-Fc in HUVECs transfected with siCTL1 or siVWF1. Average number of sprouts was quantified; a minimum 30 beads/condition/experiment were used for quantification. n=3 separate experiments. ns = not significant, **** p<0.0001, Kruskal-Wallis test. **B.** Fibrin bead assay in the presence or absence of control IgG or blocking anti-Angpt-2 antibody AZ 3.19.3 in HUVECs transfected with siCTL1 or siVWF1. Average number of sprouts was quantified; a minimum 30 beads/condition/experiment were used for quantification. n=3 separate experiments. ns = not significant; **** p<0.0001, Kruskal-Wallis test.

Tie2-Fc blocks signaling from both Angpt-1 and Angpt-2. To determine whether Angpt-2 specifically was responsible for increased sprouting of VWF-deficient HUVEC, we used an Angpt-2 functional blocking antibody (AZ 3.19.3) specifically targeting Angpt-2 but not Angpt-1^31^. HUVEC were treated with siRNA for 24 hours prior to coating onto beads, then treated with AZ 3.19.3 or IgG control from the day of embedding. Functional blockade of Angpt-2 normalised the increased sprouting of VWF-deficient cells, without affecting sprouting in control siRNA-treated cells (Figure 3B). These data demonstrated that Angpt-2 is responsible for VWF-dependent sprouting angiogenesis.

### Characterisation of ECFCs from type 3 VWD with distinct molecular defects and phenotypes

To verify the relevance of these findings in patient-derived endothelial cells, we isolated endothelial colony forming cells (ECFCs)^26, 44^ from three type 3 VWD patients with angiodysplasia (Figure 4A) and 7 healthy volunteers (Supplemental Figure 3A). VE-cadherin junctional staining confirmed the endothelial lineage of all ECFCs samples (Figure 4E,H); however, whilst VWD20B-ECFC morphology looked similar to healthy controls, VWD12B-ECFC were larger (Supplemental Figure 3B) and formed a heterogeneous monolayer. VWD21B-ECFC were also large, some showing a multinucleated phenotype, and struggled to proliferate (data not shown). Because of this, ECFCs from VWD21B were profiled for VWF but not used for further studies.

**Figure 4.**
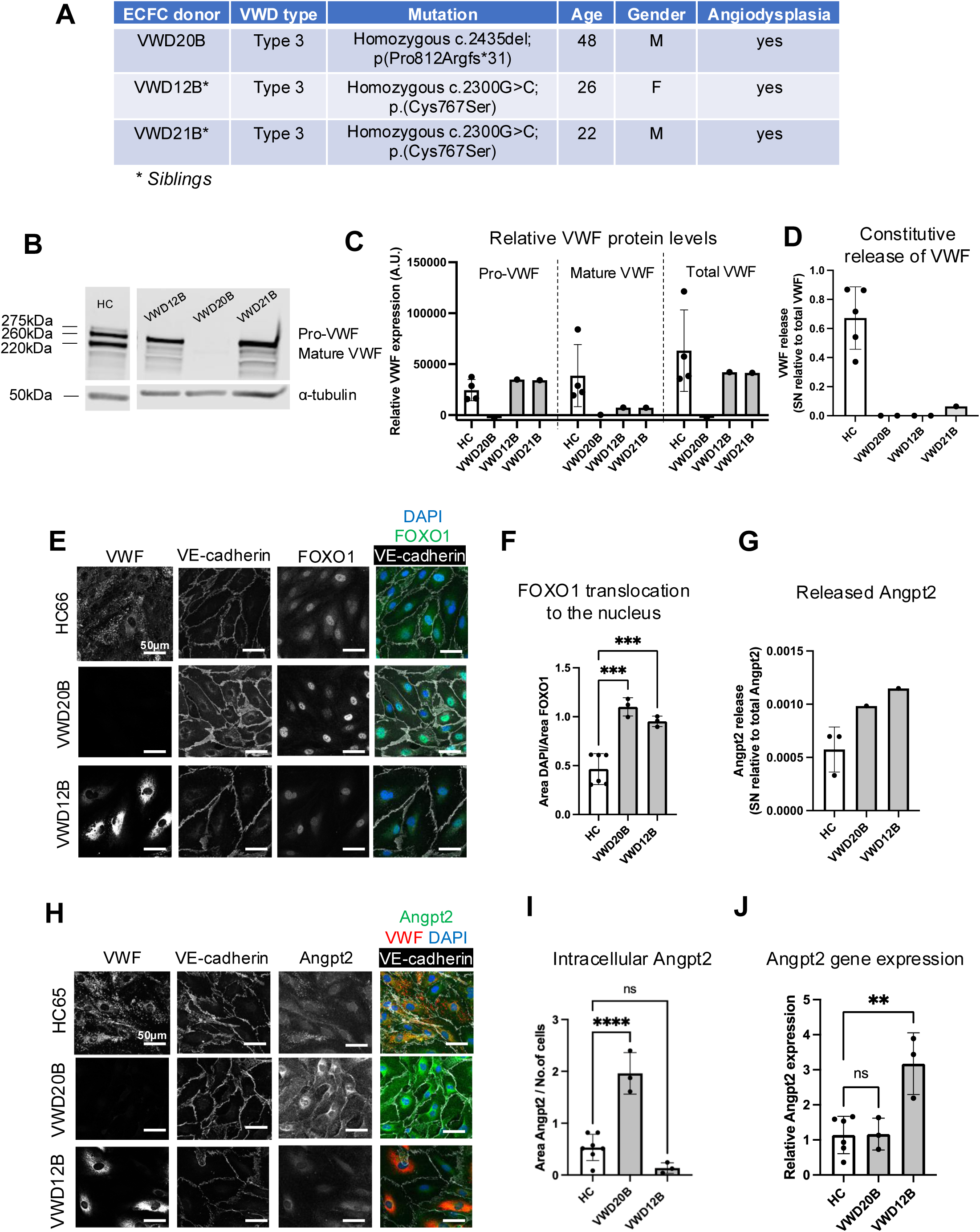
Characterisation of ECFCs from type 3 VWD and healthy controls. **A.** Table summarizing the clinical data of 3 patients with type 3 VWD. **B.** Western blot analysis of VWF protein expression in healthy control (HC) and VWD type 3 ECFC lysates. **C.** Quantification of pro-VWF, mature VWF and total VWF levels from **B**. **D.** VWF constitutive release over 48 hours from HC and VWD type 3 ECFCs. **E.** Representative images of ECFCs from HC66, VWD20B and VWD12B stained with VWF, VE-cadherin (white), FOXO1 (green) and DAPI (blue). **F.** Quantification of FOXO1 translocation to the nucleus using the ratio of DAPI/FOXO1 area. ***p<0.001, one-way ANOVA. **G.** Angpt-2 constitutive release over 48 hours from ECFCs from 3 HCs and 2 type 3 VWD patients. **H.** Representative images of ECFCs from HC and VWD type 3 stained for VWF (red), VE-cadherin (white), Angpt-2 (green) and DAPI (blue). **I.** Quantification of intracellular Angpt-2 (area/ number of cells in the field of view). n=7 images from 2 HC donors, n=3 images from VWD20B, n=3 images from VWD12B. ****p<0.0001, One-way ANOVA with multiple comparisons. **J.** Angpt-2 gene expression in ECFCs isolated from 5 healthy controls and 2 type 3 VWD patients. ** p<0.01, One-way ANOVA with multiple comparisons. All data are plotted as mean +/- S.D.

We investigated VWF levels and processing in ECFCs. Within EC, VWF undergoes complex processing, including proteolytic cleavage of its pro-peptide in the trans-Golgi network, multimerization and packaging in WPB^45^. In HC-ECFCs, Western blotting for VWF revealed 3 bands, as expected: pre-pro-VWF (275kDa), pro-VWF (260kDa) and mature VWF (220kDa). Analysis of VWF protein in ECFCs from type 3 VWD patients revealed two different VWF cellular phenotypes. VWD20B is homozygous for the most common type 3 variant in northern Europe^46, 47^, a single nucleotide deletion causing a frame shift and a premature stop codon (p(Pro812Argfs*31)). This results in a cellular null VWF defect, with undetectable levels of VWF mRNA (Supplementary Figure 3C), and undetectable intracellular (Supplemental Figure 3D) and released VWF protein (Figure 4D). In contrast, VWD12B and VWD21B (siblings) are homozygous for a point mutation at the D’D3 border of VWF (Cys767Ser), a critical region for multimerisation^48, 49^. In ECFCs from these two patients, pro-VWF is present, but no mature VWF is detectable (Figure 4B-C). VWF mRNA (Supplemental Figure 3C) and intracellular VWF protein levels are comparable to HC-ECFCs (Figure 4B-C and Supplemental Figure 3D). However, IF staining shows absence of WPBs: VWF is retained in the endoplasmic reticulum (ER) (Figure 4E,H), as confirmed by the colocalization of VWF with the ER marker calreticulin (Supplemental Figure 3E-F). Consequently, there is no VWF released from VWD12B-ECFCs and VWD21B-ECFCs (Figure 4D). The cellular phenotype of these 2 patients is reminiscent of that reported by Bowman et al in ECFCs from type 3 VWD patients with gene variants in VWF D domains^50^.

Having characterized the type 3 VWD ECFCs for their VWF defects, we tested whether the FOXO1-Angpt-2 pathway was disrupted. ECFCs from both VWD20B (null) and VWD12B (processing defect) presented increased release of Angpt-2 compared to HC ECFCs (Figure 4G) as well as increased translocation of FOXO1 to the nucleus (Figure 4E-F and Supplementary Fig 3G), in line with what observed in VWF-siRNA treated HUVEC. Similarly, Angpt-2 gene expression was increased in ECFCs from VWD12B and partly normalized by the Angpt-2 blocking antibody MEDI3617 (Supplemental Figure 3H). However, this was not observed in ECFC with the null phenotype (VWD20B) (Figure 4J and Supplemental Figure 3H). Angpt-2 intracellular protein levels were increased in the ECFC from VWD20B but not from VWD12B (Figure 4H-I). It is possible that in VWD20B ECFCs the retention of Angpt-2 in the cytoplasm might exert negative feedback towards its expression. Given the discrepancy between VWF-deficient HUVEC and VWD null ECFC, we used siRNA to inhibit VWF expression in ECFC and rule out an effect of the cellular background. In line with HUVEC, Angpt-2 mRNA was a significantly increased in siVWF-treated ECFCs and normalised by the Angpt-2 blocking antibody MEDI3617 (Supplemental Figure 3I-3J). Intriguingly, Angpt-2 blockage also decreased Angpt-2 mRNA levels in siCTL-treated ECFCs (Supplemental Figure 3I), not in HUVEC (Figure 1H and Supplemental Figure 1I). ECFCs expressed higher Angpt-2 basal levels compared to HUVEC (Supplementary Figure 3K), which may partly explain the finding.

### Angpt-2 mediates dysregulated angiogenesis in ECFCs from type 3 VWD patients

To investigate the role of VWF in angiogenesis, we developed a new 3D model of vasculogenesis/angiogenesis. Building upon our previous gut-on-chip device^32^, we enlarged the central culture channel to accommodate endothelial cells embedded into fibrin gel, flanked by two side channels for culture medium replenishment (Figure 5A-C).

**Figure 5.**
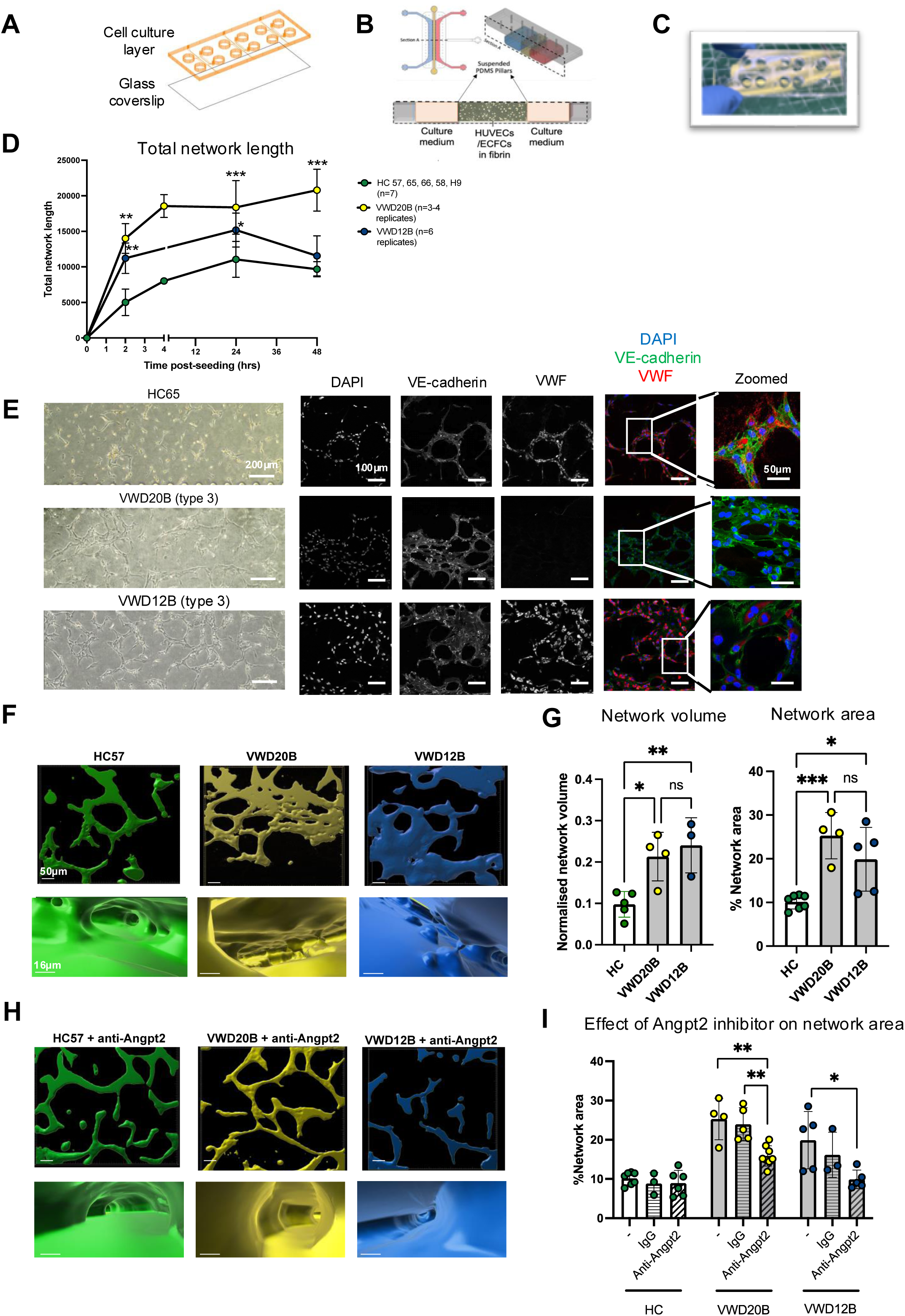
Microfluidic vasculogenesis/angiogenesis model reveals abnormal patterns of angiogenesis in ECFC from VWD type 3 patients. **A.** Schematic representation of the microfluidic devices used to develop the 3D vasculogenesis/angiogenesis model, with the cell culture layer bonded to a glass coverslip. **B.** Schematic representation of a single cell culture unit consisting of a middle channel, where endothelial cells are embedded into fibrin, and two lateral channels used to supplement the cells with culture medium. **C.** Representative picture of a microfluidic device. **D.** Quantification of total network length over time in HC (n=7 from 5 different HC donors, green), VWD12B (n=6 replicates from 1 donor, blue) and VWD20B (n=3-4 replicates from 1 donor, yellow). Data analysed at different timepoints using One-way ANOVA. *p<0.05, **p<0.01, ***p<0.001. **E.** Representative images (brightfield and confocal microscopy) of angiogenic networks formed over 48 hours using ECFCs from HCs (n=7 replicates from 5 different HC donors), VWD20B (n=4 replicates) or VWD12B (n=6 replicates). **F.** 3D rendering of angiogenic networks obtained using HC (green, n=3 different HCs) or VWD type 3 ECFCs: VWD20B (yellow) and VWD12B (blue). **G.** Quantification of network volume (normalized to image volume) and % angiogenic network area in HC or type 3 VWD ECFCs. **H.** 3D rendering of angiogenic networks from HC (green, n=3 different HCs) or VWD type 3 ECFCs: VWD20B (yellow) and VWD12B (blue) in the presence of the Angpt-2 blocking antibody MEDI3617. **I.** Quantification of % angiogenic networks area in the absence or presence of a control IgG or the anti-Angpt-2 antibody. *p<0.05, **p<0.01, One-way ANOVA with multiple comparisons. All data are plotted as mean +/- S.D.

HC-ECFCs assembled into small, organised angiogenic networks over the course of 48 hours (Supplementary Figure 4A). Compared to HC-ECFCs, ECFCs from VWD20B and VWD12B formed enhanced vascular networks with increased overall length but with wider and less remodelled structures (Figure 5D-E). A similar phenotype was observed in siRNA-transfected HC-ECFCs (Supplemental Figure 4B). Time-course analysis revealed a significant increase in total network length for VWD20B-ECFCs at both early and late timepoints compared to HC ECFCs; the same was observed for VWD12B-ECFCs at early timepoints, however these vascular networks appear to degrade after 24 hours (Figure 5D-E and Supplemental Figure 4A).

IF staining of angiogenic networks revealed the presence of punctate VWF-positive structures in HC-ECFC, reminiscent of WPB; these were absent in VWD12B-ECFCs, where VWF staining was confined to the perinuclear zone, whilst no VWF staining was detectable in VWD20B-ECFCs, in line with the 2D cultures (see Figure 4E,H).

The IF images clearly show that 2D network measurements do not capture all the abnormalities observed in VWD-ECFCs. Therefore, we carried out 3D visualization and rendering of the angiogenic structures at 48 hours (Figure 5F). These revealed remarkable, unexpected defects in VWD-ECFCs networks. Whilst HC-ECFCs formed highly remodelled vascular networks with multiple well-defined channels, suggestive of lumen structures, VWD20B and VWD12B-ECFCs organized in two poorly remodelled parallel sheets with occasional point of contact between the two, suggesting that cells attempted but mostly failed to form a lumen (Figure 5F and Supplemental Videos 1-3). This was reflected in a significant increase in network volume and network area in ECFCs from both type 3 VWD patients (Figure 5G). These data reveal an undescribed role for VWF in the control of vascular lumen formation during vasculogenesis/angiogenesis.

Finally, we tested whether these phenotypes could be normalized by inhibiting Angpt-2 activity. Brightfield microscopy showed a dose-dependent response to MEDI3617 in VWD20B-ECFCs, with no effect in HC-ECFCs (Supplemental Figure 5A-C). Remarkably, the Angpt-2 blocking antibody MEDI3617, but not the control IgG, normalised the phenotype from both VWD20B and VWD12B-ECFCs, by decreasing the network area and allowing the VWD-ECFCs to form remodelled, lumenised angiogenic structures (Figure 5H-I and Supplemental Figure 5D). To support this evidence, we repeated the experiment using HC-ECFCs transfected with siCTL2 or siVWF2. siVWF-transfected HC-ECFCs presented a significant increase in network area compared to siCTL-HC-ECFCs, which was normalized by the Angpt-2 blocking antibody MEDI3617 (Supplemental Figure 5E). These data show that increased release of Angpt-2 from VWD-ECFC is responsible for multiple angiogenic and remodeling defects that can be normalised by blocking Angpt-2 activity.

## DISCUSSION

In this study, we show that dysregulated angiogenesis in VWF-deficient HUVEC and ECFC from type 3 VWD patients can be normalised by blocking the function of the angiogenesis regulator Angpt-2. We also show that VWF modulates the synthesis of Angpt-2 in EC, via a transcriptional pathway involving the kinase AKT and the transcription factor FOXO1. These data shows that VWF controls endothelial pathways critical for vascular development, integrity and function via the regulation of Angpt-2 storage and expression. Crucially, we employ a novel microfluidic vasculogenesis/angiogenesis model which reveals defects in network formation in VWD ECFC and VWF deficient EC, pointing to unexpected functions of VWF in the control of lumen formation, opening the door to novel therapeutic strategies.

We and others have previously shown that VWF regulates the endothelial storage and release of Angpt-2^17, 23^. Angpt-2 is part of the Angiopoietins/Tie-2 pathway, a crucial system regulating vascular homeostasis and angiogenesis^51^. The current model indicates that the two ligands, Angpt-1 and Angpt-2, respectively act as an agonist and as a context-dependent antagonist of Tie-2 (rev in ^21^); thus, the relationship between these two proteins is key to vascular homeostasis^52^. Angpt-2 is more abundant at sites of vascular remodelling and in tissues where active remodelling is occurring such as ovaries, uterus and placenta ^35^. In confluent, quiescent endothelium, Tie2 activation causes activation of PI3K-AKT and phosphorylation of FOXO1^53, 54^. In our study, we show that the increased release of Angpt-2 due to loss of WPB in VWF-deficient endothelial cells promotes a positive feedback loop via AKT-FOXO1 that increases Angpt-2 expression. Crucially, we confirm imbalance of the Angpt-1/Angpt-2 *in vivo*, in the gut of VWF KO mice, where Angpt-2 levels are increased whilst Angpt-1 levels are decreased, likely due a decrease in the number of pericytes. The Visual Abstract summarises these findings and suggests indirect mechanisms of vascular instability in the absence of VWF.

Intriguingly, the Angpt-1/Angpt-2 imbalance was found only in the jejunum of VWF-deficient mice. Whether the density of vessels susceptible to angiodysplasia is higher in the jejunum is not known. In patients, angiodysplasia has been described in all regions of the gut. An imbalance in levels of Angpt-1 and -2 has been reported in the plasma of patients with small bowel angiodysplasia^55^. A large study of circulating angiogenic factors in VWD patients found a complex picture, with higher Angpt-1 levels in type 2A compared to type 2B VWD patients, and higher Angpt-2 levels in patients with increased VWF clearance^56^. Our findings suggest that increased plasma Angpt-2 levels in some VWD patients may be partly due to increased endothelial transcription of Angpt-2. Given the clinical strong correlation between GI bleeding and loss of high MW multimers, future studies will focus on the relationship between VWF multimers and the Angpt/Tie2 pathway.

During angiogenesis, Angpt-2 can signal via integrins, including αvβ3, to drive sprouting. Angpt-2 has been shown to bind to αvβ3, inducing its internalization and reducing its expression^57^. VWF can also bind αvβ3^57^ but this appears to prevent its internalization, controlling its surface levels^17^. Thus, signaling from αvβ3, which can interact with both VWF and Angpt-2 may synergize or contribute to the disrupted angiogenic phenotype of VWF-deficient cells. Another level of crosstalk may involve the VEGF/VEGFR2 system. Previous studies have suggested a role for VEGFR2 signalling in VWF-deficient EC^17^. Moreover, VWF interacts with VEGF and VEGF can both synergies with and upregulate Ang-2 expression^58–60^. The complexity of this web of ligand-receptor interactions and consequent signalling pathways requires further investigation and suggests multiple points for therapeutic intervention.

This study revealed new insight into the role of VWF in vasculogenesis/angiogenesis. Thanks to the models used here, we showed that lack of VWF results in a complex defect in sprouting, maturation and remodeling, as well as lumen formation. It is critical to point out that despite its role in vasculogenesis and angiogenesis, VWF is clearly dispensable for effective vascular development, as shown by development of animal models and importantly patients with severe deficiency of VWF, who do not present detectable defects in vascular development. This in line with the well-known redundancy of angiogenesis modulators, likely a safety mechanism to increase resilience.

The 3D vasculogenesis/angiogenesis model will serve as the basis to develop a VWD-dependent gut-on-chip model, incorporating gut epithelial cells and peristalsis^32^, to investigate the effect of the gut microenvironment on angiodysplasia. Combined with ECFCs from VWD patients, this will establish a unique personalized therapy-testing model, for the identification of new treatments for gut vascular malformations in VWD and beyond.

In conclusion, in this study we demonstrate that inhibition of Angpt-2 can normalize multiple aspects of angiogenesis in VWF-deficient endothelial cells, including ECFCs from VWD patients. Given the distribution of VWF in most vascular beds and the importance of the AKT-FOXO1 pathway in multiple vascular processes, it is likely that more VWF-dependent pathways, gene targets and functions are yet to be discovered.

## Acknowledgements

We thank Dr. Tom McKinnon, Dr. Vanessa Ho and Dr. Daisy Jones (Imperial College London) for their technical support during the project. Additionally, we thank Dr. Régis Joulia for assistance and training for using the Imaris software. We thank Ms. Litty Jose (Royal London Hospital, Barts Health NHS Trust) for assistance with patient sample collection. We would like to thank Mr. Sean Platton for the clinical lab supervision and Ms. Faith Dzumbu (Royal London Hospital, Barts Health NHS Trust) for recruiting and obtaining samples of the participants. We are grateful to AstraZeneca for providing the anti-Angpt-2 neutralising antibody AZ3.19.3. We acknowledge the FILM Facility at the National Heart and Lung Institute at Imperial College London where the confocal images were taken. The silicon wafer micropatterning was performed at PoliFAB, the micro- and nano fabrication facility of Politecnico di Milano. The production of PDMS devices was performed at the MiMic Lab of Politecnico di Milano (thanks to Alessandro Cordiale for his technical assistance) and at the Department of Bioengineering, Bionanofabrication Clean Room Core Facility at Imperial College (thanks to Dr. Florent Seichepine). This study would not be possible without the kind donation of blood for ECFC isolation from patients and healthy volunteers.

This work was co-funded by MRC (MR/X021106/1, awarded to A.M.R.), BHF (PG/16/91/32515, awarded to A.M.R. and RG/17/4/32662, awarded to A.M.R.), the NIHR Imperial Biomedical Research Centre (BRC), Rosetrees Trust (M557-F1, awarded to A.M.R.), the Imperial College–Wellcome Trust Institutional Strategic Support Fund (awarded to C.P.) and Imperial College NHLI (F26396, awarded to A.C-B.). S.Y.W. was supported by the NIHR Imperial Biomedical Research Centre (BRC).

## Authorship Contributions

A.C-B. designed and performed experiments, analysed and interpreted the data, and co-wrote the manuscript, K.S. designed and performed experiments, analysed and interpreted the data, S.Y.W. isolated ECFCs, performed experiments and analysed data, M.B. provided bioengineering input into the design and fabrication of the microfluidic devices, A.N. and J.E. performed experiments, fabricated microfluidic devices and analysed data, B.W. and D.P. performed *in vivo* experiments, A.L. and M.D. performed experiments, C.P. and K.P. provided healthy control ECFCs and technical advice, G.B. provided scientific advice, M.A.L. provided clinical expertise, S.S. recruited patients, provided patient samples and clinical expertise, M.R. provided bioengineering expertise and designed the microfluidic devices, A.M.R. conceptualised the study, secured funding, analysed and interpreted the data and wrote the manuscript. All authors reviewed the manuscript.

## Disclosure of Conflicts of Interest

n/a

## Supp. figures

**Supplemental Figure 1.**
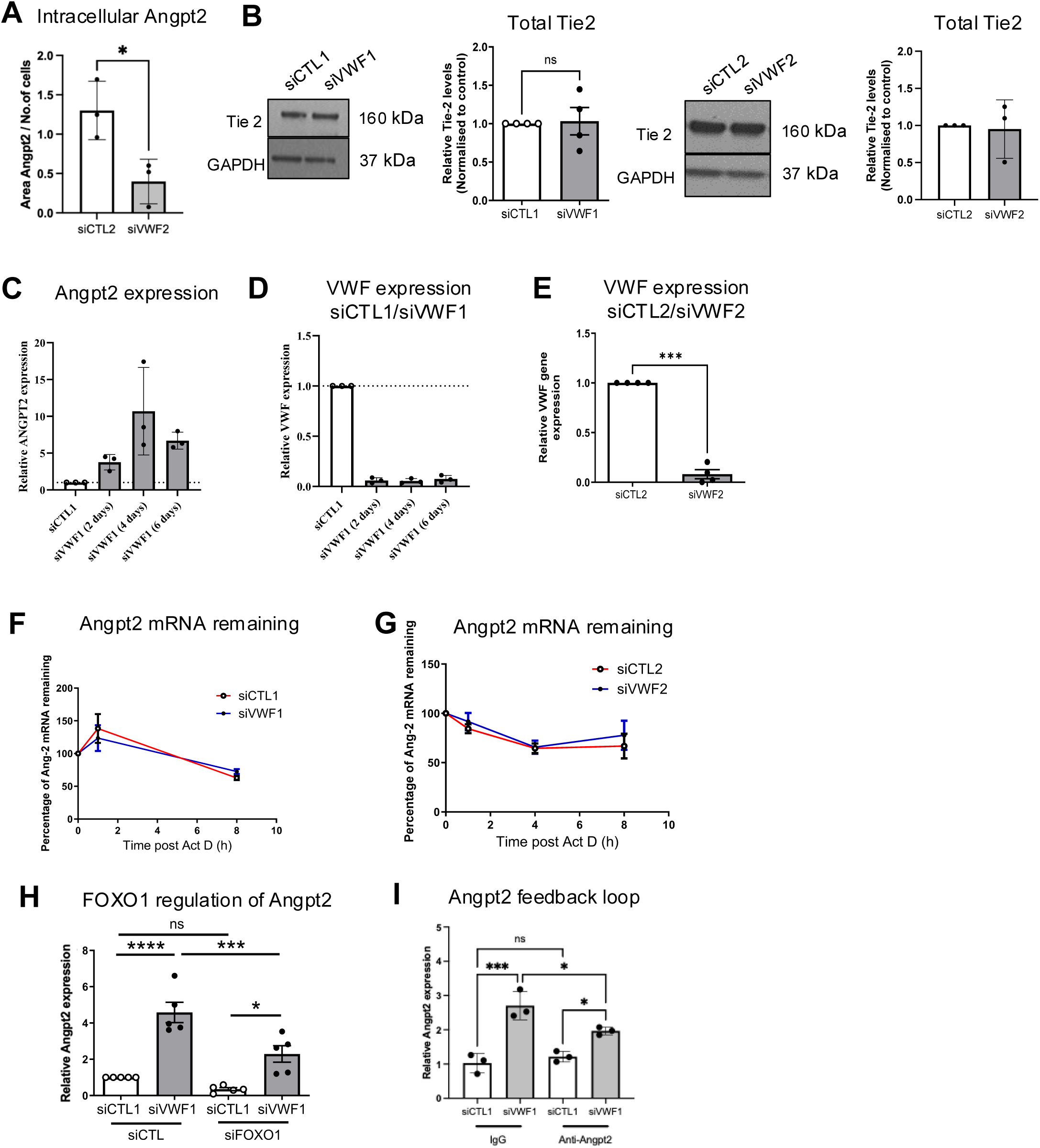

**Supplemental Figure 2.**
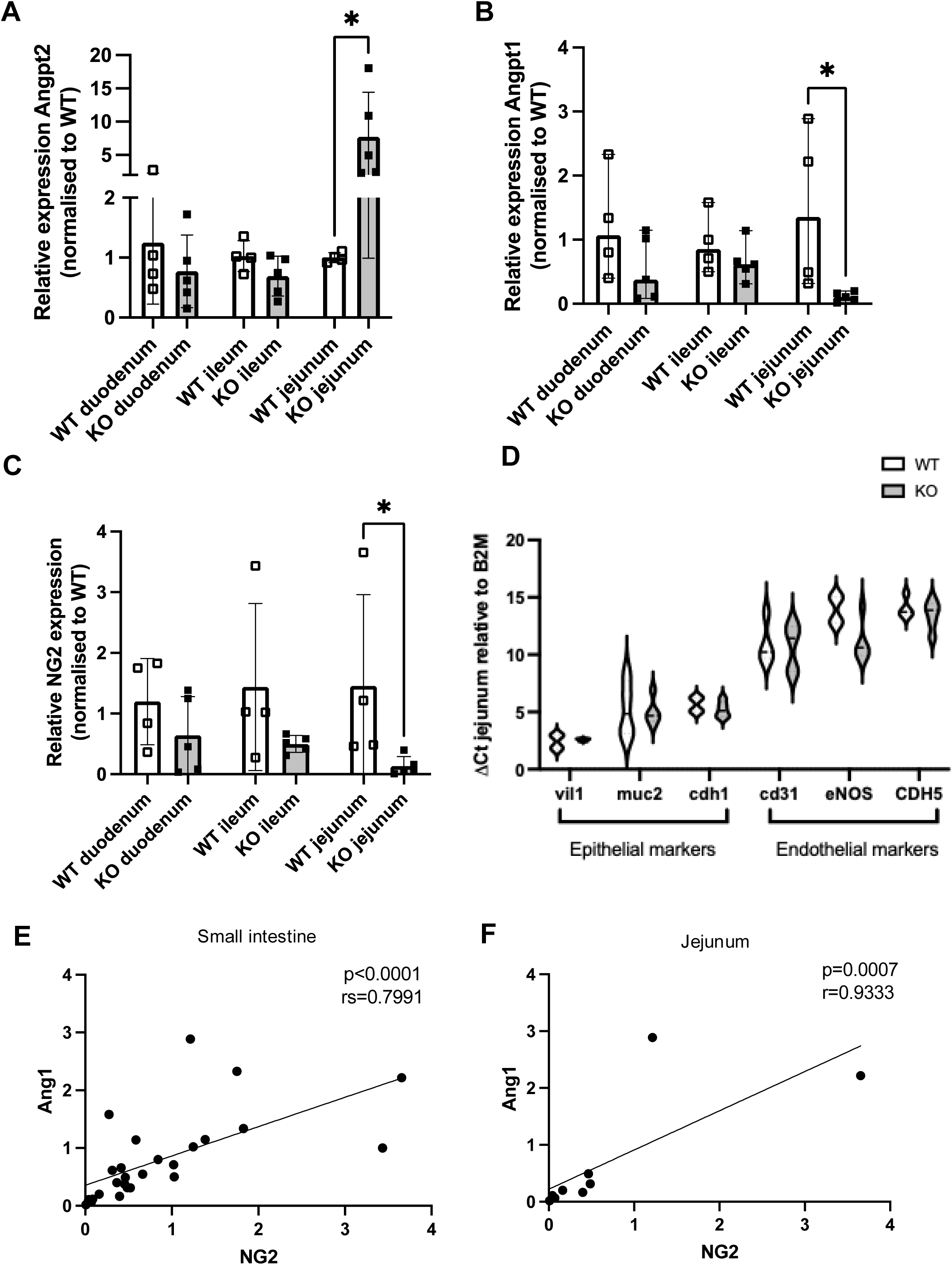

**Supplemental Figure 3.**
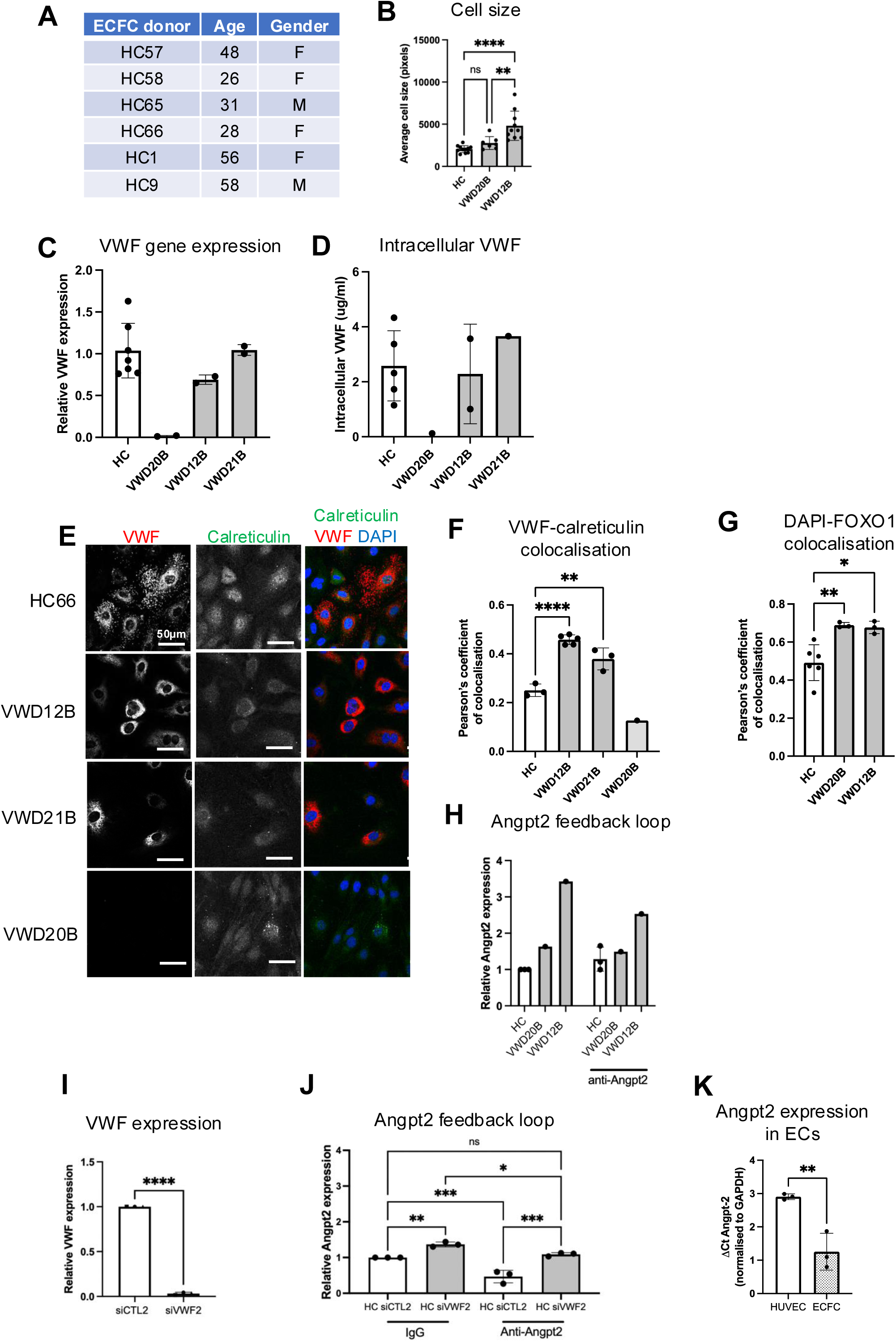

**Supplemental Figure 4.**
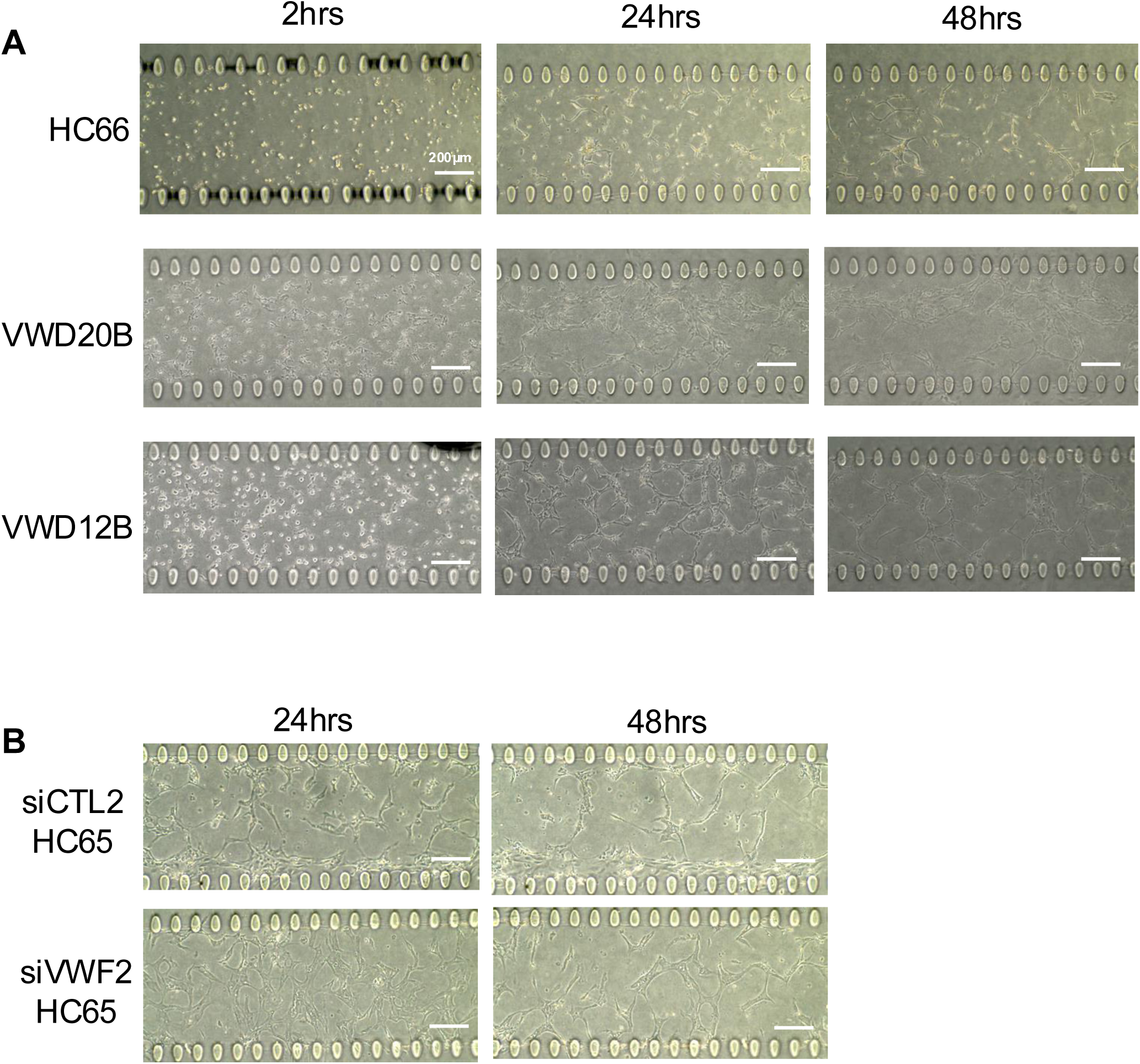

**Supplemental Figure 5.**
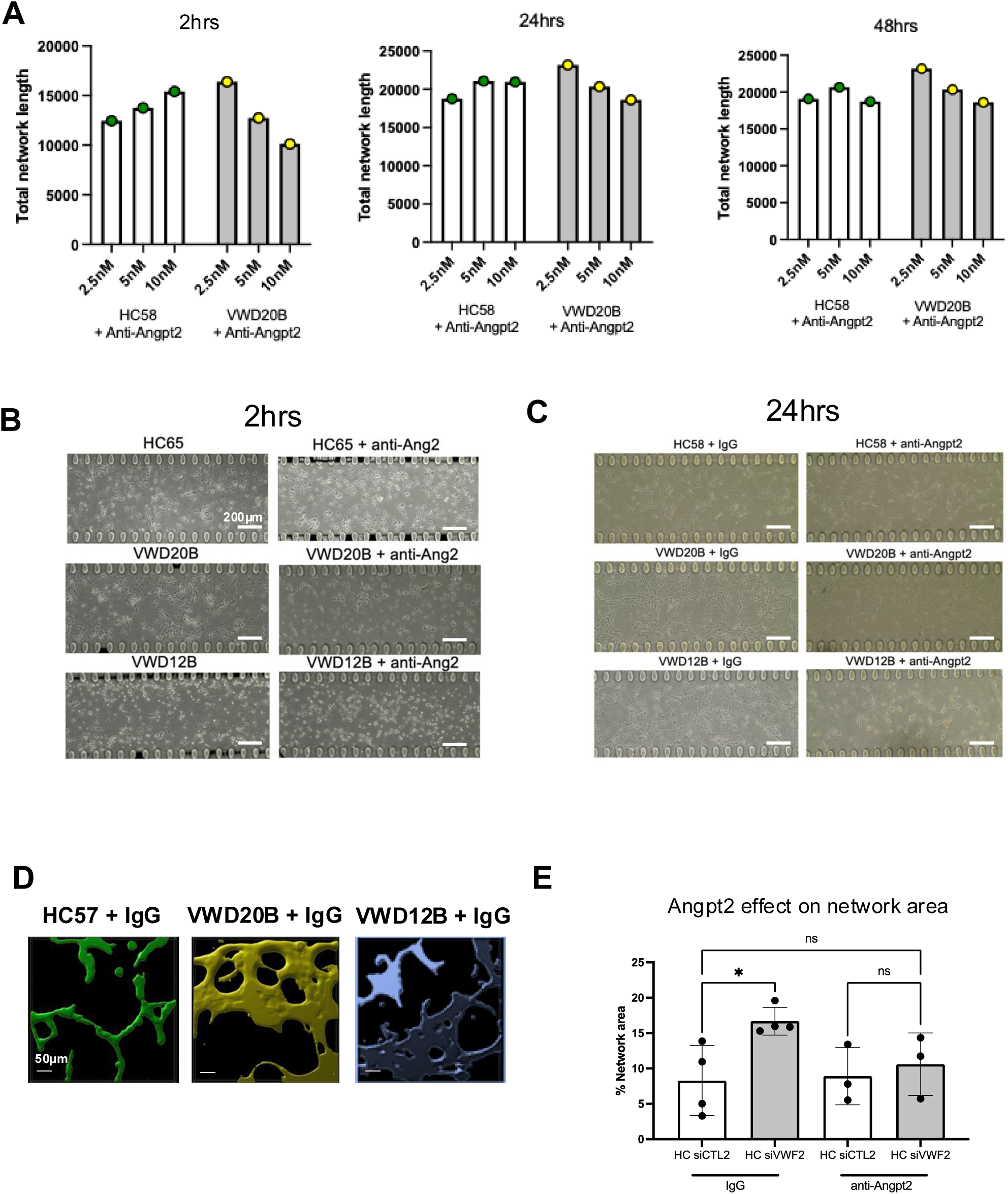

## Notes

### Competing Interest Statement

The authors have declared no competing interest.

