## Supplemental Materials for "Von Willebrand Factor Deficiency Impairs Vascular Morphogenesis via Angiopoietin-2: Relevance for Gut Angiodysplasia"

### Supplemental Methods

*Endothelial colony forming cells (ECFCs) isolation*

ECFC were isolated from blood collected in BD Vacutainer® Cell Preparation Tubes containing FicollTM HypaqueTM solution (BD Bioscience). After centrifugation the mononuclear cell layer was collected, added to PBS containing 10%HyClone serum (GE Healthcare Life science) and centrifuged again. Cells were resuspended at 3-5x$10^7$ mononuclear cells per 4ml of EGM-2 supplemented with 20% human serum (298189, Merck Life Sciences). 4ml of the mixture was added to each well of a 6-well plate pre-coated with 50µg/ml rat tail collagen type 1 (Corning).

*siRNA treatment of endothelial cells*

Two different siRNA sequences were used to target VWF sequence - siVWF1 (5-AAGGGCTCGAGTGTACCAAAA-3) from Qiagen and siVWF2 from Invitrogen (5-GGGATCTGTGATGAGAACGGA-3). In parallel All-star negative control siRNA (Qiagen), denoted as siCTL1, was used as a control for siVWF1. Silencer® Select Negative control siRNA (Invitrogen) denoted as siCTL2, was used as a control for siVWF2. These sequences have already been validated in previous studies[17]. Expression of Forkhead Box O1 (FOXO1) was inhibited using pre-validated siRNA (Qiagen) against FOXO1 (5-CCGAGTTTAGTAACAGTGCA-3)[27]; siCTL1 was used as control.

*Fibrin bead assay*

To coat the beads, cells were incubated with beads at 400 cells/bead in EGM-2 with 2%FCS. The bead-HUVEC mixture was gently shaken for 4 hours at 37°C/5% $CO_2$ and transferred into a T25 flask. The following day beads were resuspended at 500 beads/ml in 2mg/ml fibrinogen (Sigma) in PBS containing 0.15U/ml aprotinin (Sigma). 50µl of fibrinogen-bead solution was added to 24 well containing 0.625U thrombin (Sigma) and mixed gently. Mixture was allowed to solidify for 5 min at room temperature then incubated at 37°C/5% $CO_2$ for 20 min. Media was replaced every 48 hours.

*Immunoblotting*

Cells were lysed using cell lytic MT (Sigma) supplemented with protease inhibitor cocktail (Sigma), phosphatase inhibitor cocktail 2 and 3 (Sigma) and Phenylmethylsulfonlyl (PMSF) (1:100). Lysates were centrifuged for 15 min at 10 000 g at 4°C. Blots were labelled with the following primary antibodies: anti-GAPDH (1:10,000; Millipore), anti-phospho Akt S473 (1:1000; Cell Signalling), anti-pan Akt (1:1000; Cell Signalling), anti-phospho FOXO1 Thr-24 (1:1000; Cell Signalling), anti-FOXO1 (1:1000; Cell Signalling), anti-Tie 2 (1:1000; Cell Signalling). Detection was done by goat anti-rabbit IgG Dylight 680, goat anti-mouse IgG Dylight 800 (Thermoscientific) or horseradish peroxidase (HRP) linked anti-rabbit IgG (Cell Signalling) and anti-mouse IgG (GE healthcare), using Odyssey CLx imaging system (LI-COR Biosciences) and Odyssey 2.1 software. Quantification of signal was performed by densitometry and normalised against loading controls using Image J.

*Immunofluorescence 2D*

HUVEC or ECFCs were grown on 1% gelatin (HUVEC) or 50 µg/ml collagen (ECFC) coated glass coverslips in 24-well plates until confluency. HUVEC were treated as described above with siRNA to VWF or control for 48 hours. Cells were fixed using 4% paraformaldehyde (PFA) and permeabilised using 0.4% Triton X100. Staining was carried out using anti-Angpt-2 (1:500; Santa Cruz Biotechnology), anti-FOXO1 (1:200; Cell Signalling), anti-VE-cadherin (1:200; Santa Cruz Biotechnology) and anti-pTie-2 (1:200; R&D system). Nuclei were visualised using DAPI (1:500; Invitrogen). Coverslips were mounted onto glass slides using Fluoromount G™ (eBioscience). At least 3 images/coverslips were acquired using a Leica Stellaris 8 confocal microscope.

Supplemental figure legends

**Supp.Fig.1. Regulation of the Angpt-2-Tie2-FOXO1 pathway by VWF. A.** Quantification of intracellular Angpt-2 (area / number of cells in the field of view). n=3 images HUVEC transfected with siCTL2 or siVWF2. ****p<0.0001, unpaired t-test. **B.** Total Tie2 protein levels measured by Western blotting in HUVECs transfected with siCTL1/siVWF1 or siCTL2/siVWF2. **C.** Expression of Angpt-2 over time in siVWF1-treated HUVEC. **D.** Timecourse of VWF inhibition using siVWF1 in HUVECs. **E.** VWF expression in siVWF2-treated HUVEC. **F.** Angpt-2 mRNA decay after HUVECs' treatment with actinomycin D (Act D), a protein synthesis inhibitor. Angpt-2 mRNA levels are similar in siCTL1 and siVWF1-treated cells, indicating that VWF does not control Angpt-2 mRNA stability. **G.** Angpt-2 mRNA decay after HUVECs' treatment with Act D, as in panel F, with a different VWF siRNA and siCTRL pair (siCTL2 and siVWF2) -treated cells. **H**. Angpt-2 expression quantified in HUVECs transfected with siCTL1 or siVWF1, as well as siCTL or siFOXO1. **p<0.01, ***p<0.001, ****p<0.0001, One-Way ANOVA with multiple comparisons. **I.** Angpt-2 expression quantified in HUVECs transfected with siCTL1 or siVWF2, and treated with control IgG or the anti-Angpt-2 antibody MEDI3617. **p<0.01, ***p<0.001, ****p<0.0001, One-Way ANOVA with multiple comparisons. All data are plotted as mean +/- S.D.

**Supp.Fig.2. VWF$^{-/-}$ mice show an imbalance in Angpt-2 / Angpt-1 expression in the small intestine mainly driven by the jejunum. A.** Angpt-2 mRNA expression in the small intestine regions: duodenum, ileum, jejunum. **B.** Angpt-1 mRNA expression in the small intestine regions: duodenum, ileum, jejunum. **C.** NG2 mRNA expression in the small intestine regions: duodenum, ileum, jejunum. Data are plotted as mean +/- S.D. **D.** Expression of epithelial and endothelial markers in the jejunum of WT and KO mice, relative to B2M house-keeping gene. **E.** Pearson's correlation analysis between NG2 and Angpt-1 expression levels in the small intestine. **F.** Spearman correlation analysis between NG2 and Angpt-1 expression levels in the jejunum. n=4 WT, n=5 KO. Data analysed using Mann-Whitney test (*p<0.05) or Pearson's/Spearman correlation.

**Supp.Fig.3. Characterization of VWD type 3 ECFCs. A.** Table summarizing the demographic information of the healthy control ECFC donors. **B.** Average cell size (total area/no. of cells) of ECFCs from HC, VWD20B or VWD12B on coverslips imaged at 63x objective using confocal microscopy. n=11 images from 4 coverslips from 2 HC, n=7 images from 3 coverslips (VWD20B), n=10 images from 3 coverslips (VWD12B). **p<0.01, ****p<-0.0001, One-way ANOVA with multiple comparisons. **C.** VWF mRNA expression in ECFCs isolated from 5 healthy controls and 3 type 3 VWD patients. **D.** Intracellular VWF levels in 5 HC ECFCs and 3 VWD type 3 patients, measured by ELISA. **E.** Representative images obtained using confocal microscopy. Coverslips seeded with ECFCs from HC and VWD type 3 were fixed at confluency and stained for VWF (red), calreticulin (green) and DAPI (blue). **F.** Quantification of Pearson's coefficient of colocalization between VWF and calreticulin staining in panel **E. G.** Quantification of Pearson's coefficient of colocalization between FOXO1 and DAPI staining in Fig.4H. n=3 images/HC and VWD patient ECFCs. **H.** Angpt-2 expression levels in HC, VWD20B and VWD12B ECFCs, in the absence or presence of the anti-Angpt-2 antibody MEDI3617. n=3 HC donors. **I.** VWF expression in HC ECFCs transfected with siCTL2 or siVWF2. n=3 HC donors. ****p<0.0001, paired Student's t-test. **J.** Angpt-2 expression levels in siCTL2/siVWF2-treated HC ECFCs, in the absence or presence of the anti-Angpt-2 antibody MEDI3617. n=3 HC donors. **K.** Comparison of Angpt-2 expression levels in siCTL2 HUVEC vs. siCTL2 HC ECFCs, expressed as dCt of Angpt-2 normalized to GAPDH expression; n=3 repeats for HUVEC transfection, n=3 HC ECFC donors. **p<0.01, Unpaired Student's t-test. All data are plotted as mean +/- S.D.

**Supp.Fig.4. Optimisation of the microfluidic vasculogenesis/angiogenesis model to study VWF-dependent angiogenesis. A.** Representative images of networks at 2hrs, 24hrs and 48hrs obtained with brightfield microscopy using ECFCs from HC, VWD20B or VWD12B, used for quantification of network length over time in Figure 5D. **B.** siCTL2 and siVWF2-treated HC ECFCs (HC65) were embedded in fibrin and seeded in the middle channel of the microfluidic devices. Network formation was monitored using brightfield microscopy. Representative images shown at 24hrs and 48hrs timepoints.

**Supp.Fig.5. Effect of Angpt-2 inhibition on vascular network formation in VWD ECFCs. A.** Dose response of the blocking anti-Angpt-2 antibody MEDI3617 (2.5nM, 5nM, 10nM) in HC58 (green dots) or VWD20B (yellow dots) ECFCs at 4, 24 and 48 hours. n=1. **B.** Representative images of sprouting ECFCs at 2 hours post-seeding in the presence or absence of the blocking anti-Angpt-2 antibody MEDI3617. **C.** Representative images of vascular networks at 24 hours post-seeding of ECFCs, in the presence or absence of the blocking anti-Angpt-2 antibody MEDI3617 or IgG control. **D.** 3D rendering of angiogenic networks obtained using HC (green) or VWD type 3 ECFCs: VWD20B (yellow) and VWD12B (blue) in the presence of the control IgG. **E.** % angiogenic network area in HC ECFCs transfected with siCTL2 or siVWF2, in the presence of control IgG or the blocking anti-Angpt-2 antibody MEDI3617. n=3-4 images obtained with 1 HC donor (donor HC65).

**Supplemental videos. Defective vascular remodelling and lumen formation in VWD type 3 ECFCs.** Videos obtained using 3D rendering in Imaris software. **V1.** Representative video of angiogenic networks obtained with ECFCs from a healthy control (HC57), showing organized structures with successful lumen formation. **V2.** Representative video of networks obtained using ECFCs from VWD20B, showing two parallel sheets that fail to form lumenised structures and remodel. **V3.** Representative video of networks obtained using ECFCs from VWD12B, showing two parallel sheets with reduced ability to form lumenised structures and remodel.
